# Multistage classification identifies altered cortical phase- and amplitude-coupling in Multiple Sclerosis

**DOI:** 10.1101/2021.02.17.431597

**Authors:** Marcus Siems, Johannes Tünnerhoff, Ulf Ziemann, Markus Siegel

## Abstract

Distinguishing groups of subjects or experimental conditions in a high-dimensional feature space is a common goal in modern neuroimaging studies. Successful classification depends on the selection of relevant features as not every neuronal signal component or parameter is informative about the research question at hand. Here, we developed a novel unsupervised multistage analysis approach that combines dimensionality reduction, bootstrap aggregating and multivariate classification to select relevant neuronal features. We tested the approach by identifying changes of brain-wide electrophysiological coupling in Multiple Sclerosis. Multiple Sclerosis is a demyelinating disease of the central nervous system that can result in cognitive decline and physical disability. However, related changes in large-scale brain interactions remain poorly understood and corresponding non-invasive biomarkers are sparse. We thus compared brain-wide phase- and amplitude-coupling of frequency specific neuronal activity in relapsing-remitting Multiple Sclerosis patients (n = 17) and healthy controls (n = 17) using magnetoencephalography. Our analysis approach allowed us to identify systematic and non-redundant changes of both phase- and amplitude-coupling in the diseased brain. Changes included both, increased and decreased neuronal coupling in wide-spread, bilateral neuronal networks across a broad range of frequencies. These changes allowed to successfully classify patients and controls with an accuracy of 84%. Furthermore, classification confidence predicted behavioral scores of disease severity. In sum, our results unravel systematic changes of large-scale phase- and amplitude coupling in Multiple Sclerosis. Furthermore, our results establish a new analysis approach to efficiently contrast high-dimensional neuroimaging data between experimental groups or conditions.

**Highlights:**

- A novel multistage approach to analyze high-dimensional neuronal coupling data.
- Application to MEG recordings in relapsing-remitting Multiple Sclerosis patients.
- Identification of altered phase- and amplitude-coupling in Multiple Sclerosis.
- Classification of patients and controls with 84% accuracy.
- Classification confidence predicts behavioral scores of disease severity.

## 1. Introduction

Normal brain function requires coordinated interactions of neuronal ensembles that are widely distributed across the brain. Modern non-invasive brain imaging techniques, such as e.g., MEG and fMRI provide full-brain coverage, and thus, do not only allow to simultaneously measure the activity across many brain regions, but also to characterize their coupling (Hipp and Siegel, 2015; Palva et al., 2018; Wang et al., 2018), which has been linked to, for example, perception (Hipp et al., 2011), memory (Fell and Axmacher, 2011; Oswald et al., 2017; Palva et al., 2010), conscious state (Dehaene and Changeux, 2011) and various neuropsychiatric disorders (Fornito et al., 2015; Hawellek et al., 2013; Heuvel and Sporns, 2019; Kitzbichler et al., 2015; Koelewijn et al., 2017; Oswal et al., 2016; Pineda-Pardo et al., 2014; Stam, 2014).

The quantification of neuronal coupling for thousands of pairs of brain regions yields a very large, i.e. high-dimensional feature space, in which differences between populations, experimental conditions or neuronal states can be investigated. However, several caveats have to be taken into account that complicate an exploratory analysis of such high-dimensional data. First, without a hypothesis-driven restriction of the tested feature space, the direct comparison of brain-wide coupling is impeded by the large-number of connections and corresponding comparisons (Hipp et al., 2011; Pappu and Pardalos, 2014). Second, the experimental effects may entail multivariate interactions between features. These interactions are missed by a mass-univariate approach. Multiple high-dimensional classification algorithms have been developed to tease these interactions apart (see for example Kriegeskorte et al., 2008; Mur et al., 2009). However, as pointed out not all features are informative and thus classification accuracy suffers when non-informative dimensions predominate the feature space (Pappu and Pardalos, 2014). The central goal of this study was to address these problems of unsupervised feature selection to improve the comparison of brain-wide neuronal coupling between populations. To this end, we developed a novel multistage analysis approach that combines dimensionality reduction, bootstrap aggregating and multivariate classification.

To test our approach, we employed it to explore changes of brain-wide electrophysiological coupling in Multiple Sclerosis. Multiple Sclerosis (MS) is an inflammatory, demyelinating secondary neurodegenerative disease (Dendrou et al., 2015). From early disease stages on and during disease progression (Amato et al., 2010), accumulating white-matter lesions and cortical atrophy can lead to cognitive decline and physical disability (Chiaravalloti and DeLuca, 2008). The high temporal resolution of MEG and EEG allows to characterize the frequency-specific coupling of neuronal activity, which may reflect (Hipp and Siegel, 2015; Siegel et al., 2012) and even mediate (Fries, 2005) interactions in large-scale brain networks. Accordingly, electrophysiological studies in MS patients have revealed changes of neuronal coupling in specific frequency ranges (Hardmeier et al., 2012; Schoonheim et al., 2013; Sjøgård et al., 2021; Tewarie et al., 2013; Prejaas Tewarie et al., 2014). While these studies have largely focused on phase-coupling as a measure of functional connectivity, recent findings have highlighted amplitude-coupling as another mode of neuronal interactions that may provide robust connectivity information non-redundant to phase-coupling (Brookes et al., 2012; Daffertshofer et al., 2018; Hipp et al., 2012; Mostame and Sadaghiani, 2020; Siems et al., 2016; Siems and Siegel, 2020; Sjøgård et al., 2021; Wens et al., 2014). Thus, we employed our new analysis approach to resting-state MEG recordings, first, to jointly identify features of cortical phase- and amplitude-coupling that can dissociate MS patients from healthy controls, and second, to investigate if these features differ between amplitude- and phase-coupling.

Our analysis approach identified a set of principle phase- and amplitude-coupling modes that allowed to successfully classify (84% correct) RRMS patients and controls. Importantly, both coupling measures showed significant and non-redundant changes of neuronal coupling over a broad range of frequencies and cortical networks. Our results highlight non-invasive electrophysiological coupling measures as powerful new biomarkers of Multiple Sclerosis and provide a proof-of-principle for a novel approach to aid the exploration of high-dimensional coupling data.

## 2. Materials and methods

### 2.1 Subjects and dataset

We analyzed MEG data from two datasets. The first dataset was recorded at the MEG-Center Tübingen and included eyes-open resting-state MEG measurements from 34 subjects. 17 of these subjects (8 female, mean age (± std) 31.1 ± 9.6 years) were diagnosed with relapsing-remitting MS (RRMS) and 17 subjects were healthy controls (9 female, mean age (± std) 28.4 ± 4.2 years, p = 0.30). The patient group was measured prior to the first application of Tecfidera (dimethyl fumarate; BioGen Inc., Cambridge, MA, USA) with a median disease duration of 1 month (0 - 3 years interquartile range, maximum 11 years). Patients had no or mild to moderate neurological impairment, which was assessed with the Expanded Disability Status Scale (n = 16; median EDSS_total_ = 1.5, range 0 to 3.5; Kurtzke, 1983) and the Multiple Sclerosis Functional Composite (n = 13; median MSFC_totalz_ = −1.8, range −.3 to −3.3; Fischer et al., 1999). All participants gave written informed consent in accord with the Declaration of Helsinki, and the study was approved by the ethics committee of the medical faculty of the University of Tübingen.

We collected 10 minutes of eyes-open resting-state MEG data per subject. The MEG was continuously recorded with a 275-channel whole-head system (Omega 2000, CTF Systems Inc., Port Coquitlam, Canada) in a magnetically shielded room. The head position was tracked using three head localization coils fixated at the nasion and the left and right preauricular points. MEG signals were recorded with 2343.75 Hz sampling frequency and down sampled to 1000 Hz offline.

A T1-weighted sagittal MRI was obtained from each participant to construct individual high-resolution head models (MPRAGE sequence, TE = 2.18 ms, TR = 2300 ms, TI = 1100 ms, flip angle = 9°, 192 slices, voxel size = 1 × 1 × 1 mm). The subjects were scanned in a Siemens MAGNETOM Trio 3T scanner (Erlangen, Germany) with a 32-channel head coil.

The second dataset included 95 subjects from the publicly available human connectome project (HCP) S900 release (Larson-Prior et al., 2013). Participants were healthy adults in the age range between 22-35 years (n_22-25_ = 18, n_26-30_ = 40, n_31-35_ = 37). The HCP-sample included 45 females. The HCP-MEG data included three six-minute blocks of eyes-open resting-state MEG with short breaks in between measurements. Data were recorded with a whole-head Magnes 3600 scanner (4D Neuroimaging, San Diego, CA, USA) situated in a magnetically shielded room (Larson-Prior et al., 2013). Additionally, the HCP-subjects were scanned on a Siemens 3T Skyra to acquire structural T1-weighted magnetic resonance images (MRI) with 0.7mm isotropic resolution (Van Essen et al., 2013). The BTI/4D MEG system used in the HCP dataset is a potential source for dissociations to the Tübingen dataset, which was recorded with a CTF MEG system. However, we did not find any significant difference of the coupling structure (see below) between the healthy subjects measured with the two systems below 32 Hz (r_att_ > 0.95 & 0.90 for amplitude-, and phase-coupling, respectively; p_r<1,fdr_ > 0.05). For higher frequencies (>= 32 Hz) the correlation between the datasets dropped to r = 0.26 and 0.70 for amplitude- and phase-coupling, respectively, which can likely be attributed to stronger muscle artifact contamination of the HCP dataset (Fig. S1). However, importantly, while the potentially weaker reduction of muscle artifacts in the HCP-preprocessing may in principle affect component selection, it cannot lead to false-positive components for classification of patients and control subjects.

### 2.2 Data preprocessing

For the Tübingen-dataset, we first notch-filtered line noise at 50 Hz and at the first six harmonics (stop-band width: 1 Hz). Second, we visually inspected the data for muscle-, eyeblink-, and technical artifacts (SQUID-jumps). We rejected corresponding time intervals and malfunctioning or noisy channels (mean: 1 channel; range: 0 to 3 channels). Third, we high-pass filtered the data at 0.5 Hz with a 4^th^-order zero-phase Butterworth filter and split the data into two frequency bands: a low frequency band from 0.5–30 Hz and a high frequency band with frequencies above 30 Hz. For both frequency ranges, we separately performed independent component analysis (ICA; Hyvärinen and Oja, 2000). We applied the fastica algorithm with stabilization, subsequent component selection (‘deflation’), ‘pow3’ non-linearity and 1000 iterations per component. If the algorithm didn’t reach convergence within 1000 iterations the component was recomputed up to 5 times. The frequency-split approach takes advantage of the distinct spectral profile of different non-neuronal, physiological artifacts: Cardiovascular activity, eye-blinks and eye-movements most prominently show low-frequency features, whereas muscle activity is prominent at high frequencies (Hipp and Siegel, 2013). For both frequency ranges, independent components were visually inspected and artifactual components were rejected according to their topology, time course and spectrum (Chaumon et al., 2015; Hipp and Siegel, 2013). For the Tübingen dataset, from 100 extracted low-frequency components a median of 4 components was excluded (range_controls_ = 3 to 11, range_patients_ = 3 to 10). From 40 extracted high-frequency components a median of 13 components was excluded (range_controls_ = 13 to 18, range_patients_ = 8 to 21). For both frequency ranges, there was no significant difference of the number of rejected components between the two groups (both p > 0.7). After artifact rejection, the sensor-level data from both frequency bands were recombined. This recombined broad-band data was then used for all subsequent analyses.

For the HCP-dataset we used the preprocessed data as provided by the HCP pipeline (Larson-Prior et al., 2013). This included removal of noisy and malfunctioning channels, bad data segments and physiological artifacts by the iterative application of temporal and spatial independent component analysis (Larson-Prior et al., 2013; Mantini et al., 2011).

### 2.3 Physical forward model and source modeling

MEG sensors were aligned to the individual anatomy using FieldTrip (Oostenveld et al., 2010). We segmented the individual T1-weighted images and generated individual single shell boundary element method (BEM) head models. Based on these individual head models, we computed the physical forward model (Nolte, 2003) for 457 equally spaced (∼1.2 cm distance) source points spanning the cortex at 0.7 cm depth below the pial surface (Hipp et al., 2011; Hipp and Siegel, 2015; Siems and Siegel, 2020). The source shell was generated in MNI-space and non-linearly transformed to individual headspace. We co-registered the source coordinates, head model and MEG channels on the basis of the three head localization coils.

The sensor-level MEG data was projected to source space using frequency-domain linear beamforming (DICS; Gross et al., 2001; Van Veen et al., 1997). This spatial filtering approach reconstructs activity of the sources of interest with unit gain while maximally suppressing contributions from other sources.

### 2.4 Spectral analysis

We generated time-frequency estimates of the time-domain MEG signal using Morlet wavelets (Goupillaud et al., 1984). The bandwidth of the wavelets (1 spectral standard deviation) was set to 0.5 octaves with a temporal step-size of half the temporal standard deviation. We derived spectral estimates for 23 frequencies from 2.8 to 128 Hz in quarter-octave steps.

### 2.5 Neuronal coupling

We estimated neuronal amplitude- and phase-coupling by means of amplitude envelope correlations of orthogonalized signals (Hipp et al., 2012) and the weighted phase lag index (Vinck et al., 2011), respectively. Importantly, both measures are insensitive to volume conduction and might relate to distinct functional mechanisms of cortical network interactions (Daffertshofer et al., 2018; Engel et al., 2013; Siegel et al., 2012; Siems and Siegel, 2020). For the application of amplitude-coupling we used pairwise orthogonalization of the two complex signals at each time-point (Brookes et al., 2012; Hipp et al., 2012) before correlating the log-transformed power envelopes.

For both metrics all subjects and frequency bands, we generated full and symmetric correlation matrices. For further analyses, we vectorized all unique connections of these correlation matrices. We refer to the resulting vectors as coupling profiles.

### 2.6 Direct comparison of coupling

For every connection (n_c_ = 104,196) in each frequency band (n_f_ = 23), we tested for group differences in coupling strength (two-tailed Mann-Whitney U-tests) and applied false-discovery rate correction (Benjamini and Hochberg, 1995) for multiple-comparison correction within each frequency. Further, we tested which group was more likely to show stronger coupling in each frequency band by testing the distribution of the sign of significant differences (p < 0.05) against 0.5 using a binomial test. In other words, we tested the Null-hypothesis of the same probability of positive and negative signs. For the binominal statistic, we conservatively estimated the degrees of freedom at df = 40 as the rank of the forward model, i.e., the maximum amount of independently separable sources (Hipp and Siegel, 2015; Wens et al., 2015). Furthermore, we FDR-corrected the results of the binomial tests across frequencies.

### 2.7 Dimensionality reduction of coupling space

In a first step, in the HCP dataset, for each frequency and coupling measure, we applied PCA to the coupling profiles across HCP subjects (n_hcp_ = 95) in order to identify components, i.e., networks of coupling that explain most variability across subjects. Next, we projected the coupling profiles of the Tübingen patient and control subject dataset into the PCA space. For each frequency and both coupling modes, we multiplied the z-scored coupling profiles of every subject with the component eigenvectors of the PCA. In each frequency band, we used the 30 principal components with the highest eigenvalues, which resulted in 690 feature dimensions per coupling measure (coupling components).

### 2.8 Feature bagging and group classification

In a second step, we applied bootstrap aggregating (feature bagging) to identify the components that best separate between the two groups (Fig. 2A). We drew a random subset of features (n_sub_ = 10) from all 1380 features, classified the two groups using a support vector machine classifier with leave-one-out cross-validation, and repeated this procedure (n_draw_ = 2×10^7^). As control analyses, we repeated the procedure for n_sub_ = 2, 3, 5, 20, with both 5- and 10-fold cross-validation and for three other classification algorithms (decision trees, linear discriminatory analysis, naïve Bayes classifier). This procedure resulted in a distribution of classification accuracies across random feature selections. We defined the features that best separate between patients and controls by applying a threshold at the 75-percentile of this distribution. If no feature can classify between the two groups the probability of any feature to be in the subsets with the 25% highest accuracies will be equal. However, if features contain information to classify the two groups, this probability will increase. Thus, we defined the probability of a feature to be in the 25% subsets with the highest accuracies as its classification score. As control analyses, we repeated the analyses for different accuracy thresholds: 66%, 90% and 98% percentile.

**Fig. 1.**
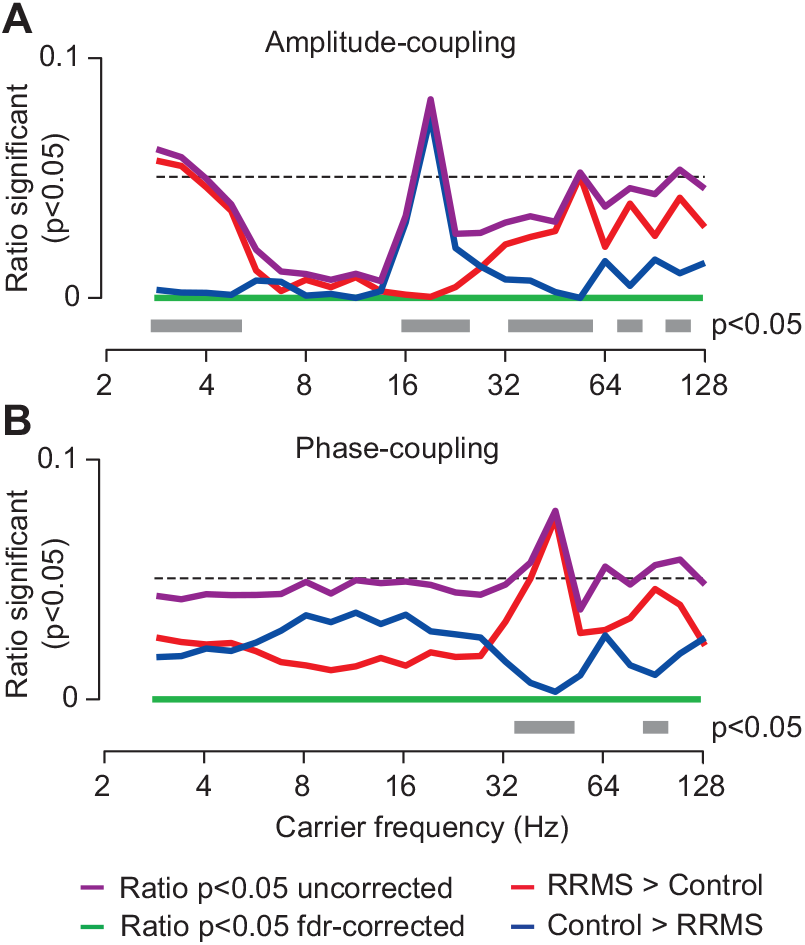
Connection-wise comparison of neuronal coupling between MS patients and control subjects. (**A**) Amplitude-coupling (orthogonalized amplitude correlations) (**B**) Phase-coupling (weighted phase-lag index). In both panels, the purple line indicates the ratio of significantly different connections (p < 0.05 uncorrected) and the green line indicates this ratio after false-discovery rate correction. The red and the blue lines show the ratio of connections with increased and decreased coupling in the patient group (rrms), respectively (p < 0.05, uncorrected). The gray bars display carrier frequencies with a significantly directed ratio of effects (p < 0.05, FDR-corrected), i.e., more connections with significantly increased or decreased coupling. The black dashed line indicates 0.05.

**Fig. 2.**
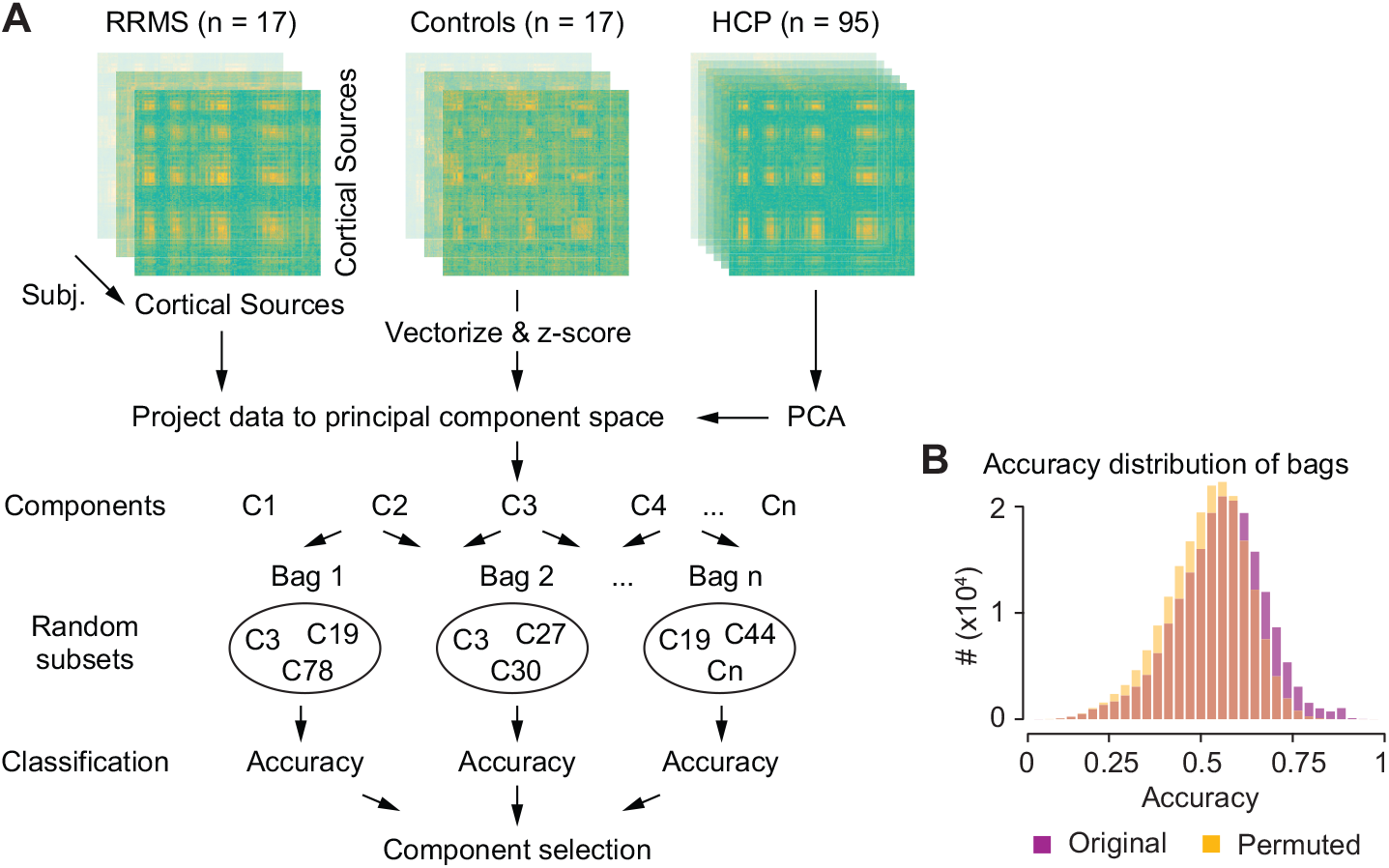
Analysis approach. For each subject, coupling measure and frequency we computed complete cortico-cortical correlation matrices and vectorized and z-scored the upper triangle. We conducted PCA on the coupling vectors of the HCP data and projected the coupling vectors of RRMS-patients and control subjects into the resulting space of principal coupling components. We applied bootstrap aggregating (feature bagging) to identify coupling components that best classify between RRMS-patients and control subjects. We drew random subsets of 10 components (bags) and, for each bag, classified (SVM) between groups with leave-one-out cross-validation. We employed a permutation statistic to select components that were more often in the top quartile of best classifying bags than expected by chance (see methods) (**B**) Distribution of classification accuracies across all 2×10^7^ bags for the original data (purple) and for randomly permuted group assignments of subjects (yellow).

Feature bagging is a central part of classic random forest classification algorithms (Breiman, 2001, 1996; Cutler et al., 2012). To elucidate the relationship of the employed approach to a standard random forest algorithm we generated a random forest classifier with feature bagging without replacement. We used 200 weak learners per learning cycle and 5-fold cross validation and repeated the computation 100 times. Subsequently, we computed the permutation importance per component and repetition. We compared the mean permutation importance over repetitions to the classification scores. Furthermore, we compared the cortical patterns of identified classifying components between the random forest approach and the proposed custom approach.

### 2.9 Statistical testing of classification scores

We used permutation statistics to test the statistical significance of every feature’s classification score, i.e., the likelihood of a given classification score under the Null-hypothesis that groups cannot be classified based on this feature. A significant feature indicates a spatio-spectral pattern of cortical coupling that separates between patient and control groups.

For each permutation (n = 1000), we randomly reassigned subjects to one of the two groups, keeping the same ratio between groups and repeated our analysis. Importantly, for each permutation, we again drew n_draw_ = 2×10^7^ feature bags per group permutation to account for the random separability of any group constellation. We then combined all classification scores (1000 permutations of 1380 feature-specific classification scores) to define one general, component-independent Null-distribution and to assign p-values for each classification score. We corrected these p-values with false-discovery rate correction across the 1380 scores (FDR-correction; Benjamini and Hochberg, 1995).

To test if groups could be significantly classified in general, we quantified how likely it was to find the identified number of significant components under the Null-hypothesis that groups could not be classified. We applied the same procedure as for the real data to every group permutation and quantified how many significant components we could identify (p < 0.05, FDR-corrected). The resulting distribution across the 1000 permutations served as the Null-distribution of significant components. All permutations had less significant features than identified for the real data (58 features). In fact, not more than one significant feature (p < 0.05, FDR-corrected) was identified for any of the 1000 permutations.

### 2.10 Classification confidence

We quantified classification confidence as a distance measure. For each subject *i*, we compared the Mahalanobis distance *D*_*mah*_ of its feature vector to the distribution of feature vectors for the patient and for the control group. We computed *D*_*mah*_ of subject i:

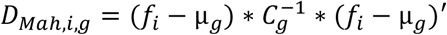

Here, *fi* describes the feature vector of each subject; *μ* and *C* describe the group *g* mean and group member covariance, respectively. We excluded the *ith* subject from the corresponding mean and covariance matrix calculation. The exponential *-1* indicates matrix inverse and the *‘* operator indicates the transpose. We then defined classification confidence as the difference between *D*_*Mah,i,con*_ *– D*_*mah,i,rms*_. Thus, a positive value indicates that a subject is closer to the patient group mean than to the control group mean and vice versa.

### 2.11 Spatial distribution of PCA components

The component coefficients derived from the HCP-dataset contain the spatial distribution of each principal component. However, the sign of these weights is inherently ambiguous. We achieved sign consistency across components by sign flipping those components for which the mean component scores within the patient group were smaller than in the control group. Hence, a positive component coefficient indicates that the coupling is relatively increased in patients whereas a negative coefficient indicates a coupling decrease.

We visualized the spatial distribution of PCA components as the average of all normalized significant components within a frequency band and coupling measure: delta (2.8 - 3.4 Hz), theta (4 - 6.7 Hz), alpha (8 - 13 Hz), beta (16 - 27 Hz), low gamma (32 - 54 Hz) and high gamma (64 - 128 Hz). For the normalization, we divided each component by the absolute 98-percentile across all its connections.

### 2.12 Unbiased accuracy estimation

To estimate the unbiased accuracy with which any new subject can be classified as patient or healthy control, we employed a second-level leave-one-out cross-validation. Leaving out each subject at a time, we applied the complete analysis pipeline outlined above, defined features spaces, and classified the left out subject.

We generated a continuous estimate of accuracy by computing the distance of the left-out subject from the decision boundary. To compare distances across different feature space realizations, i.e. left out subjects, we defined distances in units of standard deviation. We divided the distance of the left-out subject in each feature space by the standard deviation of the distances to the decision boundary of all but the left-out subjects. The sign of the distance was set positive or negative if the subject was classified as a patient or healthy control, respectively.

Finally, we separately fit a Gaussian distribution to the patient (*N*_*MS*_) and healthy control group (*N*_*con*_) distances. We derived the unbiased sensitivity, specificity and accuracy of classification from these two Gaussians:

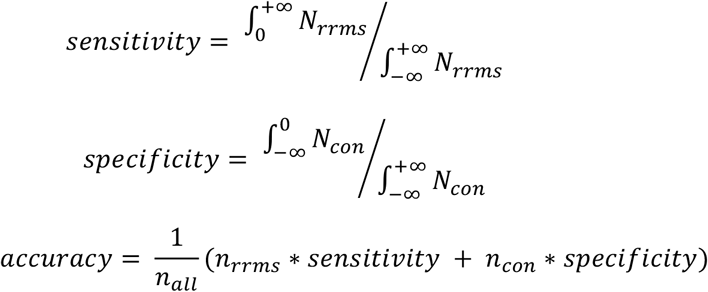

where *∫* indicates the integral and the two Gaussians, *N*_*MS*_ and *N*_*con*_, are defined by the group specific mean and standard deviation. The group sizes were defined by *n*_*MS*_, *n*_*con*_ and *n*_*all*_ for the patient group, the control group and all subjects, respectively.

## 3. Results

We compared brain-wide phase- and amplitude-coupling of frequency specific neuronal activity between 17 patients diagnosed with relapsing-remitting MS (RRMS) at an early disease stage (median EDSS = 1.5, range 0 to 3.5; Lublin and Reingold, 1996; Polman et al., 2011) and 17 healthy control subjects.

Cortical phase- and amplitude coupling was estimated from 10 min of eyes-open resting-state MEG measurements. We reconstructed cortical activity from the MEG using DICS beamforming (Gross et al., 2001; Van Veen et al., 1997) and quantified phase- and amplitude-coupling using the weighted phase-lag index (Vinck et al., 2011) and power-correlations of orthogonalized signals (Hipp et al., 2012), respectively. Both measures are insensitive to volume conduction (Brookes et al., 2012; Hipp et al., 2012; Vinck et al., 2011) and show strong intra- and inter-subject reliability (Colclough et al., 2016; Hipp et al., 2012; Siems et al., 2016; Siems and Siegel, 2020; Wens et al., 2014).

### 3.1 Direct comparison of neuronal coupling

As a first approach, we directly compared the connection- and frequency-wise coupling between patients and controls in a mass-univariate approach (Fig. 1; n_c_ = 104,196; n_f_ = 23). While differences between groups peaked around 19 Hz and 45 Hz for amplitude- and phase-coupling, respectively (Fig. 1, purple lines), these effects were not statistically significant when applying false discovery rate-correction for the number of connections tested (Fig. 1, green lines). This highlights how the high dimensionality of brain-wide coupling impedes a direct statistical comparison.

Nevertheless, the univariate comparison revealed an intriguing pattern of the sign of differences between groups (Fig. 1, red and blue lines). If there was no difference between groups, the sign of randomly significant differences in coupling (type I errors) would be equally probable in both directions. However, the observed differences deviated from this distribution for both coupling measures (Fig. 1, gray bars). For amplitude-coupling, patients showed decreased coupling in the beta frequency range (16 – 22 Hz) and increased coupling in the delta (2 – 4 Hz) and gamma (32 – 106 Hz) frequency ranges. Phase-coupling in patients was increased mainly in the low gamma frequency range (38 – 54 Hz).

In summary, although the vast number of connections impaired the direct comparison of coupling on the connection-level, the asymmetry of observed differences suggested that there are systematic and frequency-specific differences of coupling between RRMS patients and healthy control subjects.

### 3.2 Group classification based on principal coupling components

To overcome the limitations of the connection-wise analysis and to efficiently cope with the high dimensionality of the brain-wide coupling space, we next devised a multistage machine-learning approach (Fig. 2). In brief, we first identified a subspace of principle coupling components in an independent MEG dataset, then projected the patient and control data into this coupling space, and finally employed a bootstrapped classification approach to identify significant coupling differences between groups.

To efficiently reduce the dimensionality of the coupling space, we applied principal component analysis (PCA) to the z-scored brain-wide phase- and amplitude-coupling of 95 subjects from the human connectome project S900 MEG dataset (Larson-Prior et al., 2013). For each frequency and both coupling modes, we extracted the 30 principal coupling components that explained most variability of coupling across subjects (largest eigenvalues). Importantly, the identification of principal coupling components in an independent and public dataset ensured, that this identification itself was not conflated by any potential variability between patient and control groups and that the identified components could be readily applied to other research questions independent of the current dataset. We next projected the coupling profiles of patients and control subjects into the principal coupling space, which resulted in a more than 3000-fold dimensionality reduction to a total of 1380 coupling components.

We next employed multivariate classification (support vector machine, SVM) to identify significant differences of coupling between groups in the reduced coupling space. Because likely not all components are informative, classification accuracy suffers when utilizing all components at once (Pappu and Pardalos, 2014). Running the classification on the full feature space (5-fold cross-validated) could only achieve classification accuracies around chance-level: 42%, 45%, 52% and 60% accuracy for SVM, decision trees, LDA and naïve Bayes classifier, respectively. Similarly, by using every component for itself, generic interactions between components and frequencies might be missed.

We therefore employed a bootstrap approach (feature bagging; Breiman, 1996) to identify relevant components. 20-million times, we picked a random subset of 10 coupling components (without replacement) and applied SVM to classify patients and controls with leave-one-out cross-validation (Fig. 2B). For each component, we then computed a classification score as its likelihood to contribute to component-subsets with classification accuracies in the top quartile (Fig. 3A). For each component, we statistically tested its classification score using a permutation approach (p < 0.05, FDR-corrected). This procedure allowed us to identify 58 coupling components that significantly contributed to the classification of patient and control groups (Fig. 3A, p <0.05 FDR-corrected). 34 and 24 of these components were specific to amplitude-coupling and phase-coupling, respectively.

**Fig. 3.**
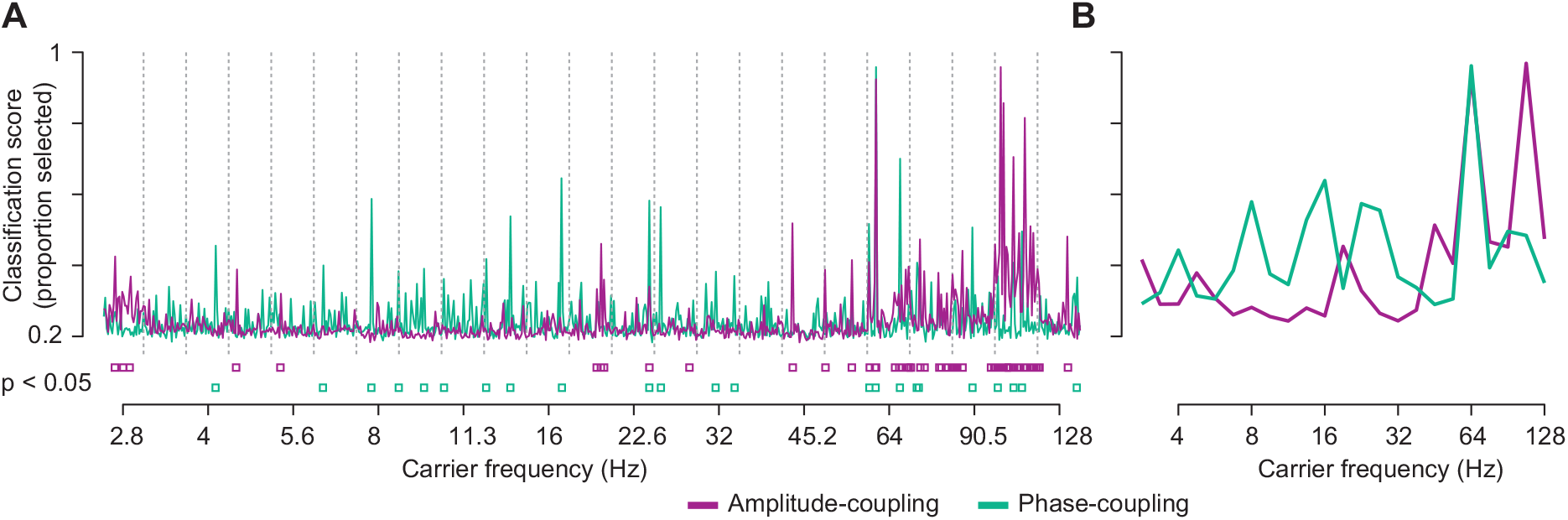
Selection of principal coupling components by classification scores. Classification scores were defined as the likelihood of each component to contribute to bags with classification accuracies in the top quartile of all 2×10^7^ bags. (**A**) Classification scores for all 30 amplitude- (purple) and phase-coupling (green) components of each carrier frequency ordered by the variance explained within each frequency. The components and eigenvalues were derived from the HCP-dataset. Dashed lines divide the carrier frequencies. Purple and green squares indicate significant classification scores for amplitude- and phase-coupling components, respectively (p < 0.05, FDR-corrected) (**B**) Maximum classification scores per carrier frequency for amplitude- (purple) and phase-coupling (green).

The employed permutation statistic also allowed us to test whether overall we could significantly classify between the two groups. We quantified how likely it was to find the identified number of significant components under the Null-hypothesis that groups could not be classified. We found that the amount of identified features was indeed highly significant (p < 0.001). Thus, RRMS patients could be significantly classified from healthy control subjects based on a specific set of principal phase- and amplitude-coupling components.

These results were robust across a broad range of parameter choices and control analyses. First, we repeated the analysis separately for amplitude- and phase-coupling components. The classification scores were similar and well correlated to those obtained when combining amplitude- and phase-coupling components (amplitude-coupling r = 0.79; phase-coupling: r = 0.65). We further repeated the analyses for different bag sizes (2, 3, 5, 20), accuracy thresholds (0.66, 0.9, 0.98), classification algorithms (naïve Bayes, decision tree and linear discriminatory analysis, see also Fig. S2) and with both 5- and 10-fold cross-validation. The results were again very similar with high correlations of classification scores between approaches (bag size: 0.92 < r < 0.96; accuracy threshold: 0.81 < r < 0.99; algorithm: 0.65 < r < 0.91; cross-validation folds: 0.93 < r < 0.98; all p < 0.05, FDR-corrected across all comparisons). Further, we found that classification scores significantly correlated with the permutation importance computed using a random forest classifier (r = 0.35, p_FDR_ < 0.05; Fig. S3).

### 3.3 Spectral and cortical distribution of altered coupling

We further examined the spectral and cortical distribution of coupling components that dissociated RRMS patients from healthy controls. For amplitude-coupling, these components were spectrally specific to low frequencies (< 5 Hz), the beta frequency range (19 Hz) and the gamma frequency range (> 40 Hz; Fig. 3B). For phase-coupling, the classification score distribution showed several peaks in the theta (4 Hz), alpha (8 Hz), beta (13 - 16 Hz & 22 - 26 Hz) and gamma bands (> 45 Hz; Fig. 3B).

To visualize the brain regions whose coupling dissociated patients from control, we first averaged the absolute coupling coefficients of all significant components for each brain region for frequencies below and above 35 Hz and both coupling measures (Fig. 4A). We split these frequency ranges because at frequencies above 35 Hz coupling estimated from MEG strongly resembles residual muscle activity (Hipp and Siegel, 2013; Siems et al., 2016).

**Fig. 4.**
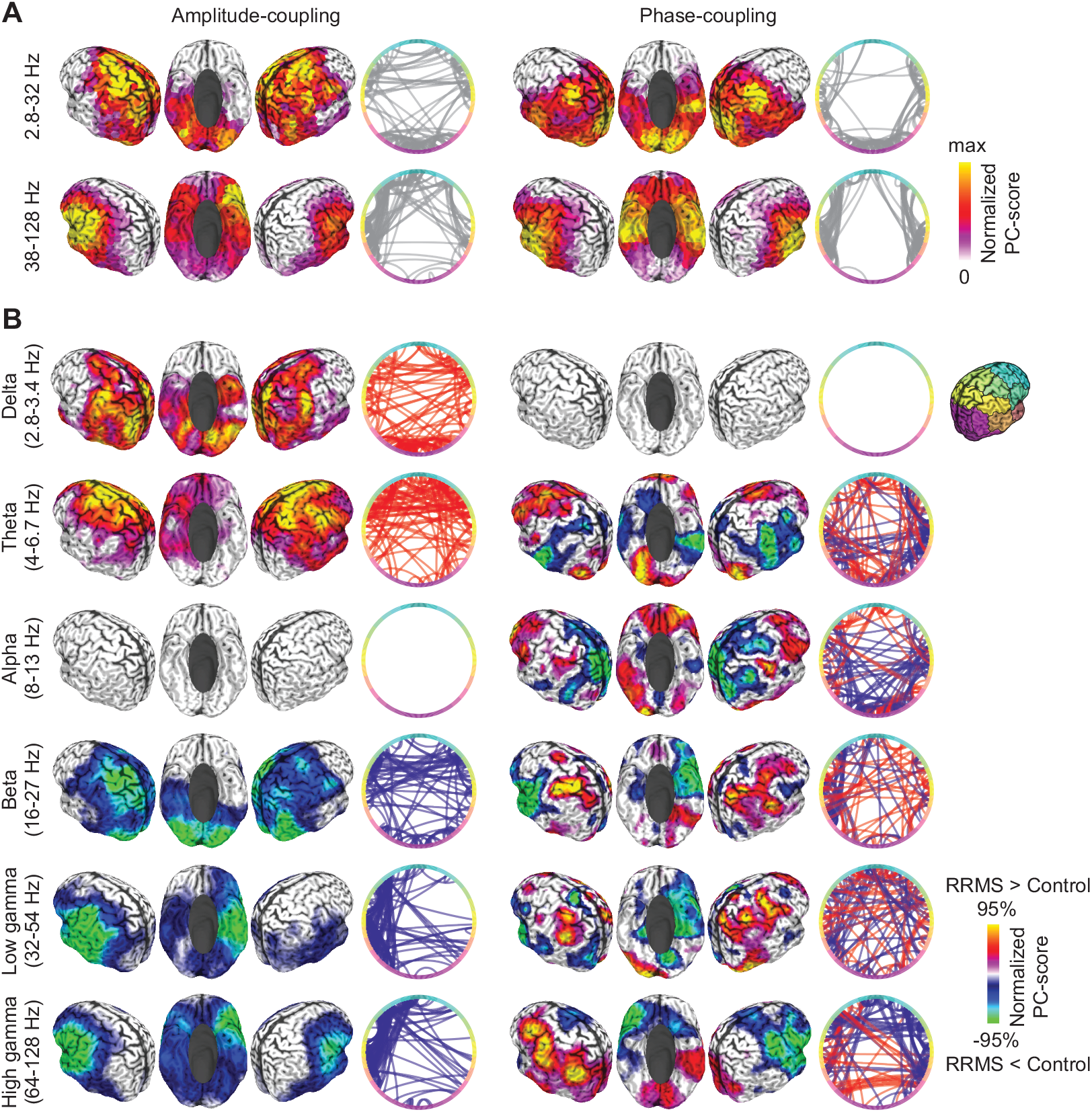
Cortical distribution of classifying coupling components. (**A**) Normalized average absolute strength of significantly classifying components for amplitude- (left) and phase-coupling (right), and for frequencies below (top) and above (bottom) 35 Hz. Colors are scaled between 0 and maximum for each panel. Circular line plots indicate connections with best classification between patients and controls (random subset of best 10%). Colors along the circle indicate cortical location, as shown on the right. (**B**) Normalized average strength of significantly classifying components for individual frequency bands and both coupling modes. Colors are scaled between the positive and negative absolute 95-percentile for each panel. Warm and cold colors indicate an increased and decreased coupling in patients, respectively. Circular line plots as in (A).

We found that, for frequencies below 35 Hz, RRMS affected intra- and interhemispheric amplitude-coupling of the medial prefrontal, dorsolateral prefrontal, pericentral, lateral parietal and extrastriate visual cortex (Fig. 4A). Phase-coupling differences in this frequency range between patients and controls peaked in bilateral pericentral, inferior temporal and medial occipito-parietal areas, mainly with altered intra-hemispheric coupling. For high frequencies above 35 Hz, for both coupling modes, we found differences between patients and controls for areas typically related to residual muscle-activity, that is bilateral anterior temporal and ventral frontal regions (Fig. 4A). These results largely overlapped with the networks found using a random forest classifier (Fig. S3). For low frequency components, the cortical patterns of the two approaches significantly correlated with r = 0.44 and r = 0.71 for amplitude- and phase-coupling, respectively. For frequencies above 35 Hz correlations were larger than 0.9 (all p < 0.05, FDR-corrected).

We next resolved the signed changes of coupling in patients within each frequency band (Fig. 4B). For amplitude-coupling, patients showed enhanced low-frequency coupling in lateral parietal and extrastriate visual areas in the delta band (2.8-3.4 Hz) and in medial and ventrolateral prefrontal cortex in the theta band (4 - 6.7 Hz). In the beta band (16-27 Hz), patients showed reduced amplitude-coupling in pericentral and visual areas. Frequencies above 32 Hz showed reduced amplitude-coupling in often muscle confounded anterior temporal and ventral prefrontal regions. Phase-coupling showed a rich pattern of changes in MS patients with typically bilateral increases and decreases of coupling within the same frequency range. Patients showed increased phase-coupling in medial prefrontal (theta, 4 - 6.7 Hz), lateral prefrontal (theta, alpha, beta & low gamma 8 - 54 Hz), as well as pericentral and lateral parietal areas (beta & gamma, 16 - 128 Hz). Phase-coupling was decreased in temporal (theta 4 - 6.7 Hz), medial and lateral parietal (alpha 8 - 13 Hz) as well anterior temporal areas (beta & gamma 16 - 128 Hz).

In summary, RRMS patients showed both, increased and decreased phase- and amplitude-coupling across a broad range of frequencies and cortical regions with highly frequency-specific changes.

### 3.4 Altered coupling predicts disease severity

If the identified changes of neuronal coupling in MS patients reflected disease-specific mechanisms, they may predict disease severity. Our results supported this hypothesis. We tested if, across patients, the confidence of classification based on neuronal coupling, i.e. how similar coupling was to that of patients as compared to controls, was correlated with two clinical measures of disease strength: the Expanded Disability Status Scale (n = 16; Kurtzke, 1983) and the Multiple Sclerosis Functional Composite (n = 13; Fischer et al., 1999). We found that indeed both clinical scores were significantly correlated with classification confidence (Fig. 5) (EDSS: r = 0.45, p = 0.04, FDR-corrected; MSFC: r = 0.49, p = 0.02, FDR-corrected). Thus, stronger coupling changes predicted a more severe disease state.

**Fig. 5.**
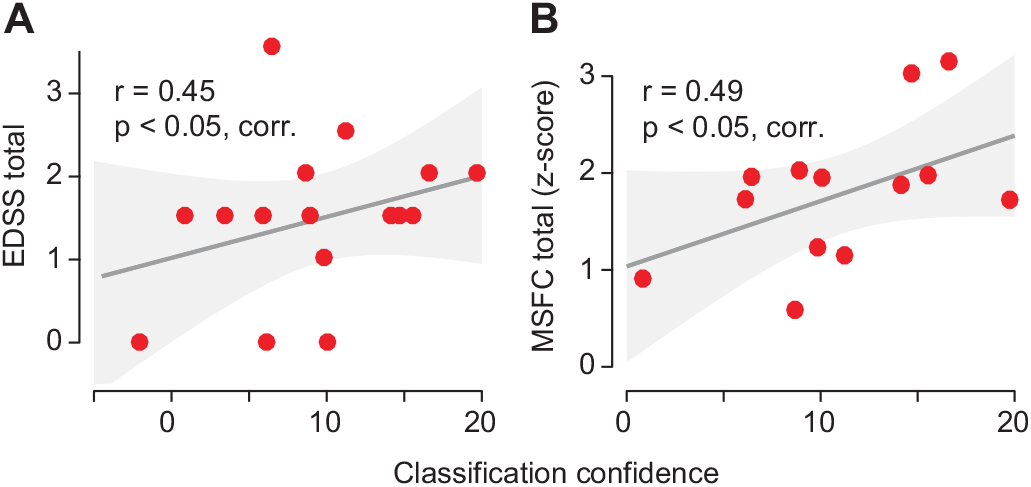
Correlations between classification confidence and behavioral scores of disease severity. Correlation between classification confidence and (**A**) the EDSS (n = 16; Expanded Disability Status Scale) and (**B**) the MSFC-scores (n = 13; Multiple Sclerosis Functional Composite). The y-axes indicate the disease severity such that an increasing value corresponds to a more severe disease state. For MSFC we mirrored the values into the positive range to fit this notation (original values MSFC_z_ = −0.3 - 3.3). Shaded areas are 95% confidence intervals of the linear model parameters.

### 3.5 Classification accuracy

In a final set of analyses, we quantitatively addressed the question how well RRMS patients could be classified based on the cortical coupling assessed with MEG. Importantly, the classification accuracy of an individual subject based on the principal coupling components identified including this subject is positively biased. Thus, to derive an unbiased estimate of classification accuracy for a novel MEG dataset, we performed a cross-validation of the entire analysis pipeline leaving out and classifying each subject at a time. We performed this analysis for all combined coupling components (n_comp_ = 1380) as well as separately for the two coupling modes (n_comp_ = 690) and frequency ranges (n_comp,low_ = 480 & n_comp,high_ = 210).

The unbiased classification accuracy using all coupling components was 84 % (chance level: 50%) and nominally higher than the accuracy obtained using any coupling mode or frequency range alone (Fig. 6A). The independent classification accuracy for amplitude-coupling (83 %) was higher than for phase-coupling (74%). Furthermore, we found that the distance to the classification boundary was only weakly correlated between amplitude- and phase-coupling (r = 0.21) (Fig. 6B). In line with the enhanced accuracy when combining both coupling modes, this is supporting evidence for non-redundant information of phase- and amplitude-coupling.

**Fig. 6.**
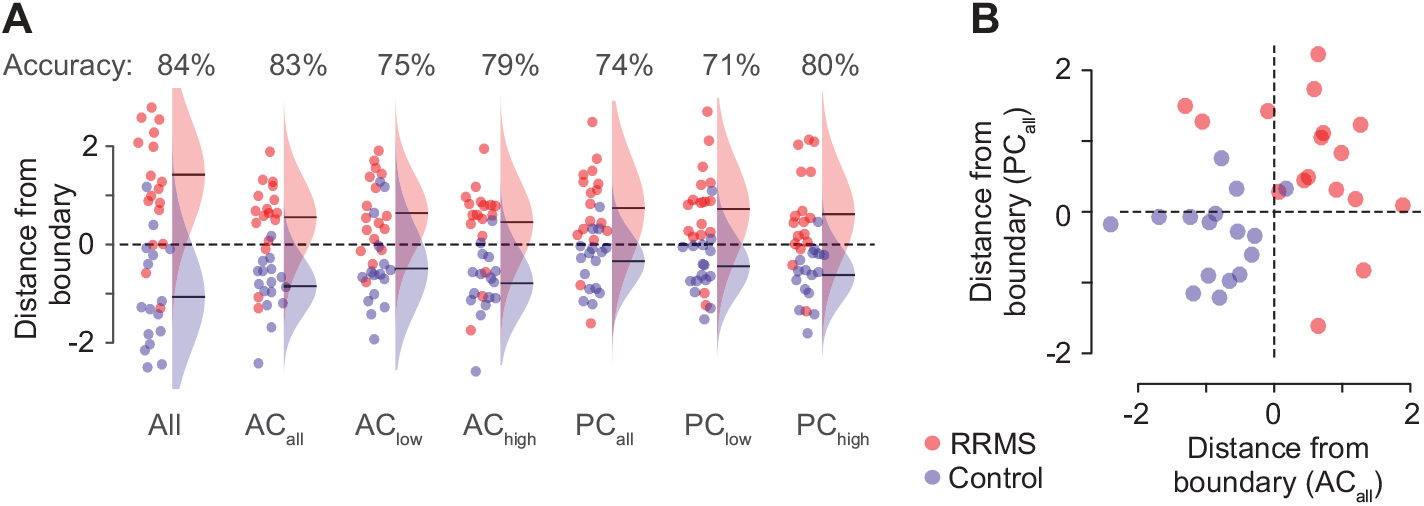
Classification accuracy. (**A**) Distance of every subject from the decision boundary (classification confidence) and Gaussian fits of the group distributions. Distances are normalized by the standard deviation across all subjects within each cross-validation fold. The circle color indicates the group; red for RRMS-patients and blue for controls. Frequencies are split at 35 Hz (low vs. high). (**B**) Each subjects’ distance from the classification boundary for all amplitude- and phase-coupling components. AC (amplitude-coupling); PC (phase-coupling).

## 4. Discussion

We devised and successfully applied a novel multistage approach to analyze high-dimensional neuronal coupling data. The approach combines dimensionality reduction, bootstrap aggregating and multivariate classification. Using this approach, we identified several brain-wide principal phase- and amplitude-coupling components that significantly dissociated RRMS patients from healthy controls with an overall accuracy of 84% and with classification confidence predicting disease severity. This study provides, to our knowledge, the first systematic comparison of brain-wide phase- and amplitude-coupling in Multiple Sclerosis patients. For both coupling modes, in RRMS patients we found both, increased and decreased coupling across a broad range of frequencies and cortical regions. At higher frequencies, effects at least partially reflected changes in muscle activity (Hipp and Siegel, 2013; Siems et al., 2016). In summary, a novel multistage classification approach revealed systematic, wide-spread and non-redundant changes of phase- and amplitude-coupling in Multiple Sclerosis.

### 4.1 Classification approach

The employed unsupervised multivariate classification approach combines several key methodological advantages. First, all analyses were carried out on reconstructed cortical activity, rather than on the raw MEG-signals. This allowed to align subjects and enhanced the signal-to-noise ratio by rejecting non-neuronal activity and focusing on signals of specific neuronal origin (Hipp and Siegel, 2013; Van Veen et al., 1997). Furthermore, we employed phase- and amplitude-coupling measures that rejected spurious coupling due to field spread.

Second, we combined dimensionality reduction and bootstrap aggregating to classify between groups in a very high-dimensional dataspace. Critically, the dimensionality reduction was based on a large and completely independent dataset. Thus, the identified principal coupling components are reflecting universal characteristics of cortical coupling across human subjects independent from the variance between the two compared groups at hand. In turn, the identified components can be readily applied to any new dataset and question at hand. We employed a bootstrap approach (feature bagging; Breiman, 1996; Tin Kam Ho, 1998) to identify several relevant dimensions in the reduced dataspace. The results were similar across a broad range of bootstrap parameters (bag size, threshold) and classification algorithms, supporting the robustness and broad applicability of this approach. The presented approach allows to directly integrate further classification methods. Multivariate distance measures such as cross-validated Mahalanobis distance (Allefeld and Haynes, 2014) and multivariate regression analyses are other possible extensions of the presented pipeline. With feature bagging as a central component, the employed approach bears similarity to classic random forest classifiers. Indeed, we found that the key results were compatible with those of a standard random forest algorithm.

Third, we employed cross-validation at two critical stages of the analysis pipeline. First, we cross-validated the classification between groups for every bootstrap aggregate of coupling components and combined this permutation statistics. This prevented overfitting for the identification of classifying coupling components. Second, we cross-validated the classification accuracy of the entire analysis pipeline, which ensured an unbiased estimate of the classification accuracy for new datasets.

Overall, high dimensional feature spaces are a common problem in brain-connectivity analyses and biomedical research in general. Classification of groups of samples on the entire space is often not feasible because of uninformative dimensions and classification on individual dimensions may be computationally impractical, might miss multivariate interactions, and is difficult to control for multiple comparisons (Hipp and Siegel, 2015; Pappu and Pardalos, 2014; Wang et al., 2018). In the present case, this is well illustrated by the failure to detect significant coupling changes in a direct connection-level comparison. In these situations, the analyses approach employed here provides a flexible tool that can be readily adapted to explore a high-dimensional space.

Despite these benefits and the validity of the present approach, it should be noted that the comparatively small sample-size of the present dataset (n = 34) limits generalizability. External validation within a larger and independent sample is necessary to further improve parameters of the analysis pipeline and to establish generalizability of the identified coupling changes.

### 4.2 Wide-spread bidirectional changes of cortical coupling

Our results provide new insights into the spatial and spectral distribution of cortical coupling changes during RRMS. We found altered coupling across the entire investigated frequency range (2.8 to 128 Hz). This adds to a growing but heterogenous body of studies that have identified MS-related changes of rhythmic neural activity or coupling across various different frequency bands (Cover et al., 2006; Figueroa-Vargas et al., 2020; Hardmeier et al., 2012; Schoonheim et al., 2015, 2013; Schoonhoven et al., 2019; P. Tewarie et al., 2014; Tewarie et al., 2013; Van Schependom et al., 2014). For phase-coupling, our approach revealed both increases and decreases within the same frequency range. This may have contributed to the heterogeneity of previous findings and highlights the advantage of separating cortical networks at the source-level.

Rhythmic coupling at different frequencies reflects interactions in specific neuronal micro- and macro-circuits (Donner and Siegel, 2011; Siegel et al., 2012). The spectrally widespread nature of our findings suggests that MS leads to alterations across many different circuit interactions. This may entail not only local and large-scale cortico-cortical interactions, but also cortico-subcortical interactions. For example, the altered phase-coupling in the alpha frequency range found here may reflect altered cortico-thalamic interactions associated with MS-related thalamic atrophy (Schoonhoven et al., 2019; Tewarie et al., 2013).

The observed broad spatial distribution of altered coupling across the cortex accords well with its broad spectral distribution. Changes involved both, local and large-scale coupling. In general, the affected cortical networks resembled the known association with specific frequency ranges, such as altered theta- or beta-coupling in midfrontal and sensorimotor regions, respectively. This suggests that RRMS alters the strength of interactions in common brain networks associated with specific frequencies. It remains to be determined, to what extent also the frequency of these interactions is affected (Schoonhoven et al., 2019).

At frequencies above 35 Hz, RRMS patients showed decreased frontotemporal coupling, which well resembled the reported distribution of residual muscle activity in EEG and MEG (Hipp and Siegel, 2013; Siems et al., 2016). Thus, the decreased high-frequency coupling may be due to decreased muscle activity, which may in turn result from both, the disease itself as well as from motivational differences between patients and control subjects.

Motivational or cognitive differences may generally contribute to classification results between patient and control groups. In this context, the finding that classification confidence predicts disease severity within the patient group is particularly important. This finding suggests that the identified changes indeed reflect disease specific effects rather than general motivational differences between patient and control groups.

Several different mechanisms may contribute to such disease specific changes. On the one hand, lesions of white-matter tracts may directly cause a reduced coupling of the connected neuronal populations. On the other hand, such reduced anatomical connectivity and functional coupling may induce a number of indirect effects. For example, the decoupled populations may be part of a larger neuronal network. Local decoupling in this network may lead to a global decrease of coupling, compensatory enhancement of coupling or a mere shift of the network dynamics. Similarly, changes within one network may again lead to both, decreases as well as increases of coupling in other brain networks (Fornito et al., 2015; Helekar et al., 2010; Heuvel and Sporns, 2019; Schoonheim et al., 2015; Stam, 2014). Importantly, all these changes could span a broad range of temporal-scales, which may even lead to opposite immediate and long-term effects of the disease.

The interplay of all these mechanisms may explain the wide-spread, complex and often bi-directional pattern of coupling changes observed here. The present results set the stage for future studies to disentangle the different mechanisms underlying these changes. For this, longitudinal investigations will be particularly important (Schoonheim et al., 2015).

### 4.3 Coupling mode specific changes

Amplitude correlation of orthogonalized signals has recently been introduced as a robust and spectrally specific marker of cortical coupling (Hipp et al., 2012; Siems et al., 2016; Siems and Siegel, 2020). Consistent with other recent evidence (Sjøgård et al., 2021), our results show that cortical amplitude-coupling is systematically altered in RRMS patients already at an early disease stage. Moreover, our findings uncover markedly distinct changes for phase- and amplitude-coupling. There were more coupling components that dissociated the groups and higher classification accuracy for amplitude-coupling than for phase-coupling (amplitude/phase-coupling: 34/24 components; 83/74% accuracy). This suggests that coupling changes are more robust for amplitude-coupling.

Furthermore, while phase-coupling showed effects in all but the delta-band, amplitude-coupling was altered in all but the alpha-band. The effects were also spatially dissociated. Below 35 Hz, amplitude-coupling showed strongest changes in medial and lateral prefrontal cortex as well as in pericentral and medial parietal areas, while phase-coupling showed strongest effects in pericentral, medial occipitoparietal and inferior temporal cortex. Additionally, while amplitude-coupling showed either consistent increases or decreases of coupling within each band, phase-coupling showed bidirectional effects within each band. For frequencies above 35 Hz, amplitude-coupling showed higher sensitivity to residual muscle activity. Finally, we found that the two coupling modes showed different sensitivities to different subgroups of subjects.

Overall, our results show that phase- and amplitude-coupling are sensitive to at least partially distinct changes of cortical coupling in MS. Accordingly, while both coupling modes could independently dissociate patients from healthy controls, the combination of coupling modes increased classification accuracy. Our findings suggest that amplitude-coupling provides a robust biomarker of changes in large-scale network dynamics that may be synergistically combined with phase-coupling measures. Adding other features of population activity could further improve classification performance. In particular, source-level power may provide additional information. Further studies are required to test if power in specific frequency bands and cortical regions shares variance with the identified diagnostic connectivity structure. This may be because of power directly affecting connectivity as well as because of signal-to-noise ratio fluctuations (Pesaran et al., 2018; Siems and Siegel, 2020).

In sum, the observed differences between coupling modes add to converging evidence that at least partially distinct neuronal mechanisms underlie amplitude- and phase-coupling (Daffertshofer et al., 2018; Engel et al., 2013; Siegel et al., 2012; Siems and Siegel, 2020).

### 4.4 Summary and conclusions

In summary, we devised a new multistage analysis approach that combines dimensionality reduction and bootstrapped multivariate classification to identify disease-related neuronal coupling changes. Our approach can be readily adapted to other scientific questions, and thus, holds potential for the comparison of experimental populations or conditions in high-dimensional data spaces.

Explorative application of our approach on a comparatively small sample of patients and healthy controls uncovered systematic changes of large-scale cortical phase- and amplitude-coupling at an early disease stage of Multiple Sclerosis. Changes were coupling-mode specific and included decreases as well as increases across wide-spread frequency ranges and cortical networks. Our results highlight the potential of non-invasive measures of neuronal phase- and amplitude-coupling as powerful biomarkers for brain-network disorders.

## Data availability

The HCP data are available for download from https://www.humanconnectome.org/. The Tübingen dataset and code are available from the authors upon reasonable request.

## Acknowledgements

We thank Hannah Krämer, Yeho Kim and Gabi Walker-Dietrich for help with data acquisition.

## Funding

This work was supported by the European Research Council (ERC) StG335880 (M.S) and the Centre for Integrative Neuroscience (DFG, EXC 307) (M.S.). The authors acknowledge support by the state of Baden-Württemberg through bwHPC, the German Research Foundation (DFG) through grant no INST 39/963-1 FUGG (bwForCluster NEMO), and a grant from Biogen Idec GmbH (U.Z.).

## Competing interests

The authors declare no competing financial interests.

## CRediT authorship contribution statement

**Marcus Siems:** Conceptualization, Methodology, Investigation, Formal analysis, Software, Writing - original draft, Writing - review & editing. **Johannes Tünnerhoff:** Conceptualization, Investigation, Resources, Writing - review & editing. **Ulf Ziemann:** Conceptualization, Resources, Writing - review & editing, Funding acquisition. **Markus Siegel:** Conceptualization, Methodology, Resources, Writing - original draft, Writing - review & editing, Supervision, Funding acquisition.

## Supplementary Material

### Supplementary Figures

**Fig. S1.**
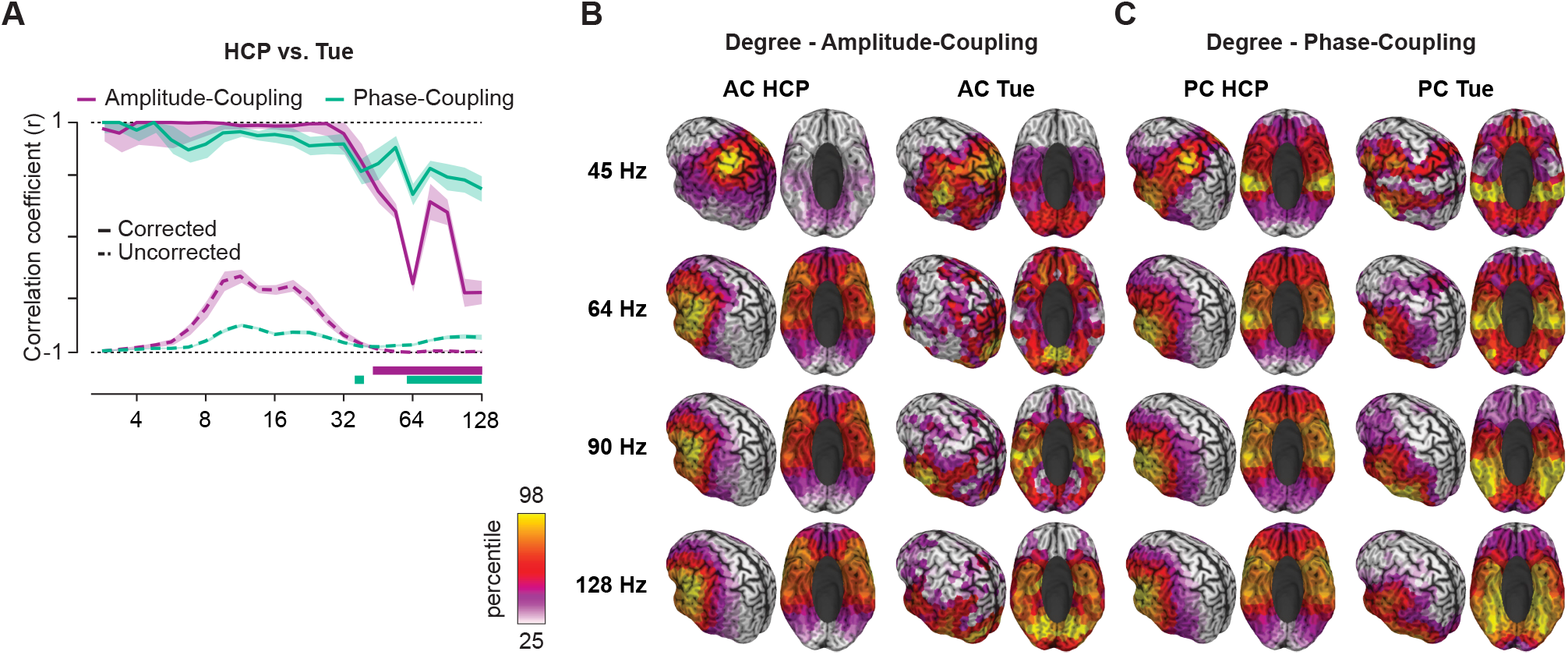
Comparison of coupling between HCP and Tübingen control subjects datasets. (**A**) Raw (dashed lines) and attenuation corrected correlation (solid lines) between connectivity profiles of the HCP (BTI) and the Tübingen (CTF) dataset (healthy controls only). Coupling is compared both for amplitude- (purple line) and phase-coupling (green line). Shaded areas indicate the standard error of the mean over leave-one-out pseudo-values. Colored bars indicate significant inter-site correlation below 1 (r_corrected_ < 1, p < 0.05, FDR-corrected). (**B**) & (**C**) Mean coupling patterns (Degree) for the high carrier frequencies (45 – 128 Hz) for (B) amplitude- & (C) phase-coupling generated for the HCP (left) and Tübingen control subjects (right). The color scale is set to between the 25^th^ to 98^th^ percentile within each panel.

**Fig. S2.**
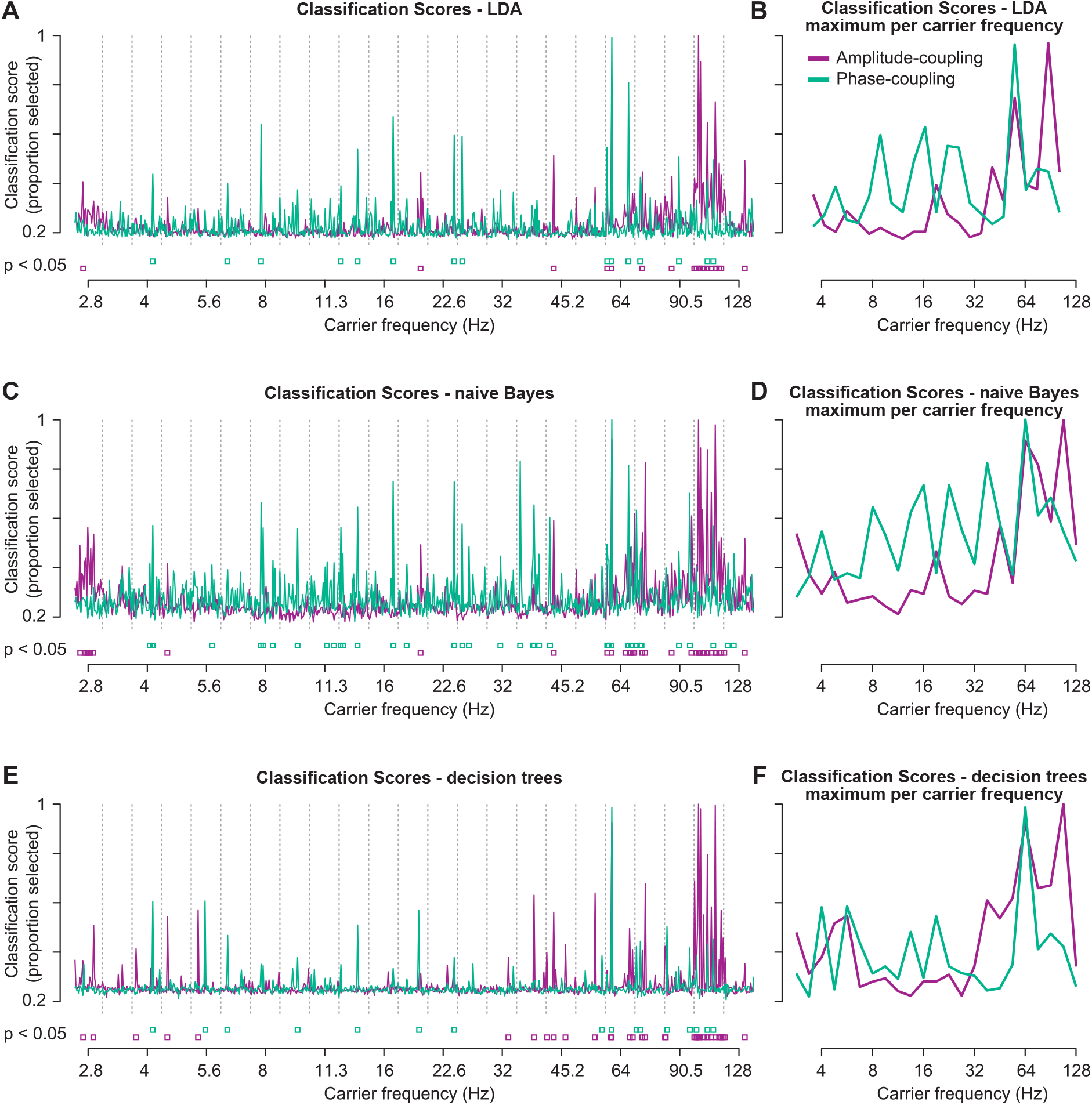
Comparison between different classification methods. (**A, C, E**) Classification scores for all 30 amplitude- (purple) and phase-coupling (green) components of each carrier frequency for LDA, naïve Bayes, and decision tree classifiers. Purple and green squares indicate significant classification scores for amplitude- and phase-coupling components, respectively (p < 0.05, FDR-corrected). (**B, D, F**) Maximum classification scores per carrier frequency for amplitude- (purple) and phase-coupling (green) for LDA, naïve Bayes, and decision tree classifiers. All results are shown as in Fig. 3 for SVM.

**Fig. S3.**
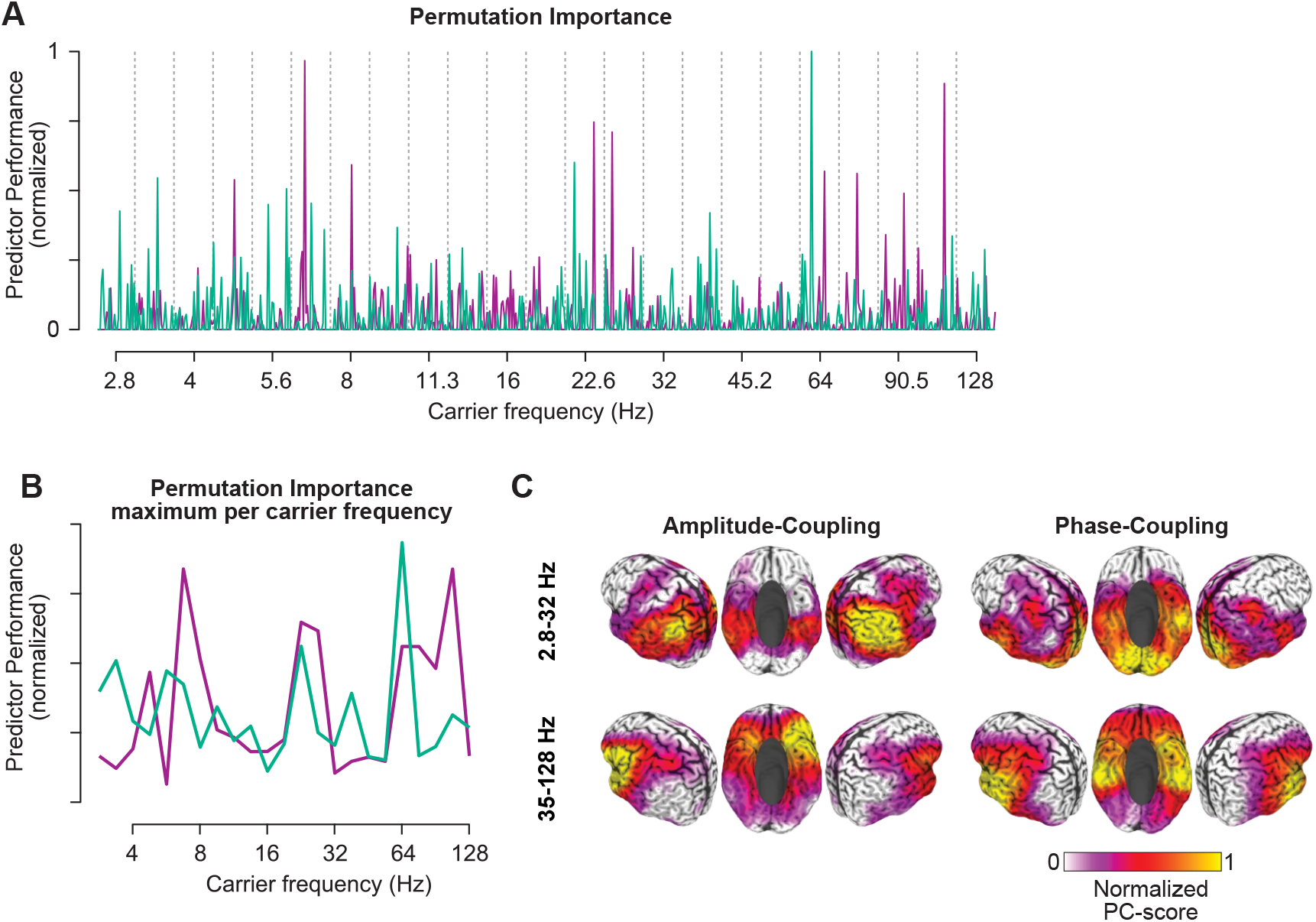
Classification results from a random forest classifier. (**A**) Weight per component (Permutation Importance) for all 30 amplitude- (purple) and phase-coupling (green) components of each carrier frequency for the classification between patients and control subjects. (**B**) Maximum permutation importance per carrier frequency for amplitude- (purple) and phase-coupling (green). (**C**) Normalized average absolute strength for amplitude- (left) and phase-coupling (right), and for frequencies below (top) and above (bottom) 35 Hz. All components were normalized and scaled according to their respective predictor importance prior to averaging. Colors are scaled between 0 and maximum for each panel. All results are shown as in Fig. 3 & 4 for SVM.

## Notes

### Competing Interest Statement

The authors have declared no competing interest.

### Summary of Updates

The reviewers' feedback has led us to carefully revise the entire manuscript and to perform several new analyses, with interesting new results that we included in the revised manuscript. The most prominent changes and new results are: 1)We performed a new set of analyses systematically comparing coupling profiles between the HCP and Tuebingen dataset to exclude scanner-related differences (new Supplementary Figure and text-section 2.1). 2)We performed a new analysis using a standard Random-Forest classifier to elucidate the relationship of our presented analysis pipeline to related existing approaches. We found qualitatively compatible results and included the new analysis in the revised manuscript (Supplementary Figure 3 and text sections 2.8, 3.2, 3.3, and discussion) 3)We re-computed the feature bagging step in our analysis pipeline with different cross-validation variants (5-fold & 10-fold), which demonstrates the robustness of our findings. We included the results in the revised manuscript. 4)We performed a new analysis directly applying classification without the feature bagging step. The results highlight the benefit of feature bagging and are included in the revised manuscript. 5)We included the results of different classification algorithms in a new Supplementary Figure 2 (compatible to Fig. 3 of the original manuscript).

